# ROS in tobacco stigma exudate affect pollen proteome and provoke membrane hyperpolarization

**DOI:** 10.1101/2020.04.28.065706

**Authors:** Breygina Maria, Klimenko Ekaterina, Shilov Eugeny, Mamaeva Anna, Zgoda Viktor, Fesenko Igor

## Abstract

1.

ROS are known to be accumulated in stigmas of different species and can possibly perform different functions in plant reproduction. Here we confirm the assumption that they affect pollen by altering ion transport through the plasma membrane; as a more deferred effect, pollen proteome is modified. We detected ROS in stigma exudate, found hyperpolarization in exudate-treated growing pollen tubes and used flow cytometry of pollen protoplasts to compare the effects of fresh exudate and exogenous H_2_O_2_ on pollen tube plasmalemma. Exudate causes plasmalemma hyperpolarization similar to the one provoked by H_2_O_2_, which is abolished by catalase treatment and ROS quencher MnTMPP. Inhibitory analysis indicates the participation of Ca^2+^- and K^+^-conducting channels in the observed hyperpolarization, linking obtained data with previous patch-clamp studies *in vitro*. For a deeper understanding of pollen response to ROS we analyzed proteome alterations in H_2_O_2_-treated pollen grains. We found 50 unique proteins and 20 differently accumulated proteins that are mainly involved in cell metabolism, energetics, protein synthesis and folding. Thus, pollen is getting ready for effective resource usage, construction of cellular components and rapid growth.

**Highlights:**

- The active substance in stigma exudate is H_2_O_2_
- H_2_O_2_ causes hyperpolarization mediated by the activation of cation channels.
- H_2_O_2_ affects pollen proteome; we found 50 unique proteins.

## 2. Introduction

Recently the idea of reactive oxygen species (ROS) as physiological regulators in plants has been widely accepted (Mittler *et al*. 2011; Kangasjärvi & Kangasjärvi 2014; Gilroy *et al*. 2014; Mangano *et al*. 2016) and supported by multiple studies; their role in plant reproduction has been studied and discussed (McInnis *et al*. 2006a; Hiscock *et al*. 2007) but comprehensive information about it was still missing.

ROS are known to be accumulated in stigmas of different species (McInnis *et al*. 2006a; Hiscock *et al*. 2007). In olive O_2_^•-^ and NO are generated by stigma papillae (Zafra *et al*. 2010). The studies of ROS dynamics revealed species-specific patterns of ROS accumulation in species with wet or dry stigmas and self-incompatible plants (Zafra *et al*. 2010, 2016). In all species studied ROS were present on stigma awaiting pollination. The dynamics of antioxidant enzyme expression (SOD, peroxidase) as well as corresponding enzymatic activity studied in sunflower is thought to contribute to complex ROS dynamics on stigma (Sharma & Bhatla 2013). Significant complexity is the identification of particular ROS. In one of the studies mentioned above, selective H_2_O_2_ quencher sodium pyruvate (NaPyr), histochemical stain TMB, relatively selective to H_2_O_2_ and NO-selective dye DAF-2DA were used for this purpose (McInnis *et al*. 2006b). It was reported that the main ROS produced by the stigma papillae was H_2_O_2_. Pollen, however, is capable of lowering ROS level on stigma due to antioxidant properties of exine (Smirnova *et al*. 2012) and/or the reducing capacity of pollen diffusate (žárský *et al*. 1987), thus, ROS level regulation is carried out by joint efforts of pollen and stigma.

Since the discovery of ROS in the receptive stigma, hypotheses about their functions have been put forward (Hiscock *et al*. 2007): (1) primary recognition of pollen of their own species, (2) activation and control of pollen grain germination, (3) pathogen defense. For (1) experimental evidence has not been obtained yet. It seems much more likely that molecules of higher complexity are involved in recognizing the pollen of its own species (Rejón *et al*. 2014). Regarding hypothesis (3), pathogens carried with pollen are of particular interest: a similar transmission mechanism is known for some fungi, many viruses and viroids (Mink 1993; Agrios 2005; Card *et al*. 2007; Kawamura *et al*. 2014). The indirect evidence given in favor of (3) is that ROS accumulation is also characteristic for nectar and tissues secreting it (Carter & Thornburg 2004): it is known that nectaries produce viscous nutritious secret (as do stigmas) and thus can serve as “gates of infection” for various phytopathogens (Agrios 2005). Thus, the hypothesis on the protective function of ROS on stigma does not yet have reliable evidence, although there is no reason to reject it. Most likely, the accumulation of ROS on the receptive stigma surface is multifunctional: at the same time, ROS provide metabolism activation in rehydrated pollen grains and protect progeny from pathogen transmission.

Hypothesis (2) has convincing evidence obtained using simplified *in vitro* model system (Potocký *et al*. 2007; Smirnova *et al*. 2009a): ROS quenching with Mn-TMPP (100–300 µM) significantly lowers the growth rate (Potocký *et al*. 2007). Adding moderate concentrations of Mn-TMPP (H_2_O_2_ and O_2_^-**•**^ quencher) as well as DPI suppressed pollen germination both in tobacco and spruce (Smirnova *et al*. 2009a; Maksimov *et al*. 2018). On the other hand, the addition of exogenous ROS in moderate concentrations (100 µM for tobacco) activates germination (Smirnova *et al*. 2009a); Despite this rather convincing data, detailed studies and comparison of pollen grain physiology in the presence of both ROS and the stigma exudate were needed to clarify the function of ROS on the receptive stigma surface.

*In vitro* studies have revealed the link between ROS and ion homeostasis in pollen indicated by the sensitivity of the ion transport systems to H_2_O_2_. Thus, in lily pollen the sensitivity of Ca^2+^- and K^+^-conducting channels to H_2_O_2_ was demonstrated by patch-clamp (Breygina *et al*. 2016). In tobacco pollen protoplasts H_2_O_2_ caused membrane hyperpolarization and increased [Ca^2+^]_cyt_ (Maksimov, Breygina, *et al*. 2016). The sensitivity of Ca^2+^ fluxes to peroxide has also been demonstrated on spheroplasts from pear pollen tubes (Wu *et al*. 2010). In growing lily pollen tubes H_2_O_2_ caused shifts of membrane potential (MP) gradient, pH gradient and [Ca^2+^]_cyt_ increase (Podolyan *et al*. 2019). Thus, targets for H_2_O_2_ have been found in male gametophyte, but to what extent the application of H_2_O_2_ reflects the effect of exudate itself?

In this study we detected ROS in fresh stigma exudate, used flow cytometry of pollen protoplasts to compare the effects of stigma exudate and exogenous H_2_O_2_ on pollen tube plasmalemma and to test the hypothesis on the involvement of Ca^2+^- and K^+^-conducting channels in this effect. We used both pollen tubes and pollen protoplasts and different detection methods to ensure the effect of exudate on membrane potential. For a deeper understanding of pollen response to ROS we analyzed proteome alterations in H_2_O_2_-treated pollen grains.

Pollen germination and early tube growth strictly depends on protein synthesis that is relatively independent of transcription (Mascarenhas 1993), so proteomics identification of proteins differentially expressed in germinating pollen will generate important molecular information. Proteomic studies from *Arabidopsis* (Holmes-Davis *et al*. 2005; Noir *et al*. 2005) and rice (Dai *et al*. 2006) have revealed that mature pollen presynthesizes a complement of proteins required for pollen function but a significant difference between proteome in mature and germinating pollen has been revealed (Zou *et al*. 2009). Functional category analysis indicated that these differentially expressed proteins are mainly involved in signaling, cellular structure, transport, defense/stress responses, transcription, metabolism and energy production. This finding indicated that proteomic data are essential to our understanding of pollen function.

In the present study, we wanted to address the difference between proteomes of germinated pollen in control and H_2_O_2_-treated suspensions, which would highlight proteins involved in physiological processes activated by ROS.

## 3. Materials and methods

### 3.1 Plant cultivation, stigma exudate collection and protoplast isolation

Plants of *Nicotiana tabacum* L. var. Petit Havana SR1 were grown in a climatic chamber under controlled conditions (25°C, 16h light) in vermiculite. The plants were watered with salt solution (Nitsch 1965). Anthers were removed just before flower opening and dried at 25°C for three days. The pollen was collected and preserved at –20°C.

Aliquots of the dried pollen were incubated in a moist chamber at 25°C for 2 h, germinated, and used for protoplast isolation. The pollen grains were germinated in Petri dishes in standard medium at 25°C for 1 h. The pollen tubes were centrifuged (1 min, 3300 g), transferred to a hypertonic medium containing enzymes, and incubated at 29°C for 1 h. The protoplast suspension was washed to remove the enzymes and cell wall debris (two cycles of centrifugation at 600 × g for 30 s and resuspension in hypertonic medium free of the enzymes). The process of protoplast formation is shown in Fig.1a.

For exudate collection stamens were removed on the eve of flower opening. The next day pistils with wet (perceptive) stigmas were removed from open flowers and soaked in PBS pH 7.4 for 1 h for exudate washout (8 pistils in 200 µl) (Cresti *et al*. 1986). The solution (“mentioned as exudate”) was used within 2 hours.

### 3.2 Cultivation medium

The standard medium was 0.3 M sucrose, 1.6 mM H_3_BO_3_, 3 mM Ca(NO_3_)_2_, 0.8 mM MgSO_4_, and 1 mM KNO_3_ in 25 mM MES_Tris buffer, pH 5.8. The hypertonic medium for protoplast isolation was the same as the standard medium with an additional 0.4 M sucrose, 0.7 M mannitol, 2% cellulase from *Trichoderma viride*, and 1% pectinase from *Aspergillus niger* (Maksimov, Breigin, *et al*. 2016).

### 3.3 Spectrofluorometry

Deesterified form of DCFH was used to detect ROS in stigma exudate. Deesterification of DCFH-DA stock solution (10 mM) was performed during 1 h with 10x volume of 10mM NaOH right before the experiment and finished by the addition of equal volume of PBS pH 6.0 (Smirnova *et al*. 2009a). Exudate was stained for 20 min in the dark with 10 µM DCFH and measured with RF-5301 (Shimadzu) spectrofluorometer. Fluorescence was excited with Xe lamp at 445 nm, measured at 500-550 nm.

### 3.4 Membrane potential mapping in pollen tubes

Mapping of MP in pollen tubes was performed as described earlier (Maksimov *et al*. 2018) with Di-4-ANEPPS (Sigma) which allows to detect accurately spatiotemporal variations of MP. Pollen tubes were stained with 2 μM Di-4-ANEPPS and immediately used for microscopy. Red fluorescence (>590 nm) was excited in two channels: blue (450–490 nm, F_b_) and green (540–552 nm, F_g_), F_b_/ F_g_ ratio is inversely proportional to MP value (higher ratio = depolarization).

### 3.5 Flow cytometry of pollen protoplasts

Flow cytometry was performed on protoplasts from tobacco pollen tubes. Intact pollen grains are covered with a thick wall which has autofluorescence and is nonspecifically stained with fluorescent dyes, thus inconvenient for use in flow cytometry. Protoplasts are formed form apical parts of pollen tubes (Fig. 1a) and don’t contain nuclei (Maksimov, Breygina, *et al*. 2016). They are usually considered as a model system for checking the action of substances on the plasma membrane of pollen vegetative cell.

**Figure 1.**
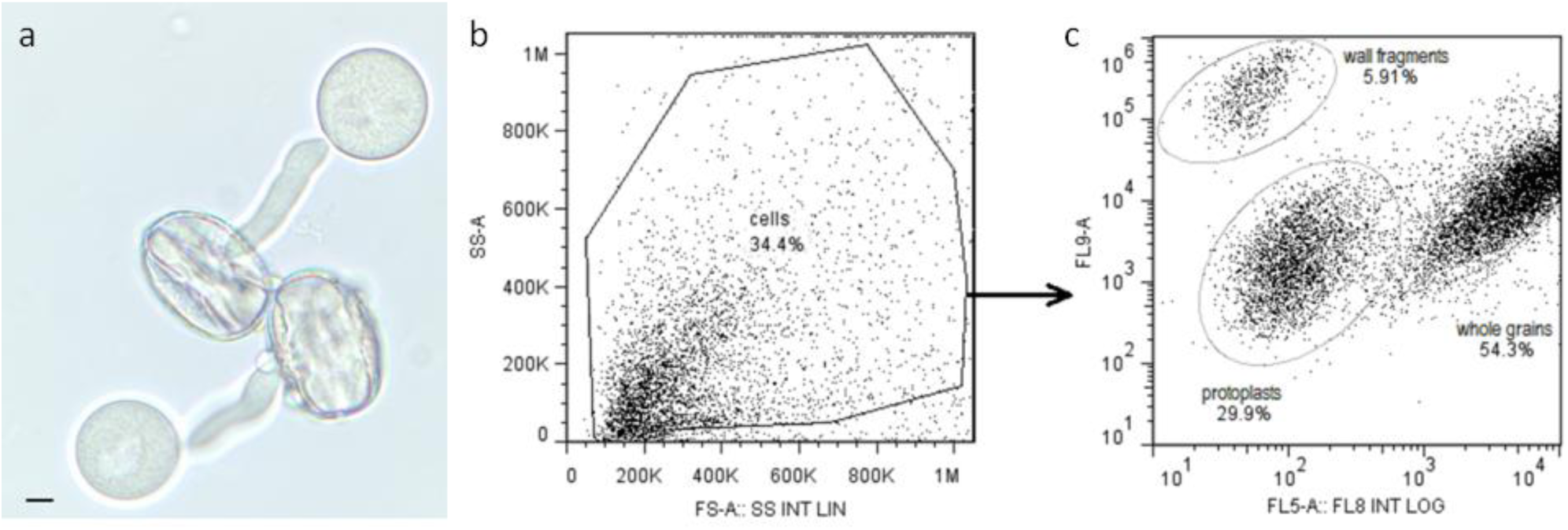
Flow cytometry of tobacco pollen protoplasts. a – protoplast formation from pollen tubes in enzymatic solution; b,c – gating strategy based on (b) forward and side scatter, (c) tinopal fluorescence (channel FL9, excitation 405 nm, detection 450/40 nm band pass) vs. far red autofluorescence (channel FL5, excitation 488 nm, detection long pass 755 nm).

MP of pollen protoplasts was assessed by staining with 1 µM DiBAC_4_(3) for 5 min (Breygina *et al*. 2010) on Gallios flow cytometer (Beckman Coulter), analysis was performed with FlowJo software (TreeStar). Tinopal staining (1 mg/l) was used to distinguish between the population of protoplasts and cell debris, pollen walls and intact grains. This population was the only one with fluorescence changing in the presence of voltage-affecting agents, for example, NaN_3_ (we observed a noticeable increase in signal as depolarization occurred). The population gating strategy based on a) forward and side scatter, b) tinopal fluorescence (channel FL9, excitation 405 nm, detection 450/40 nm band pass) vs. far red autofluorescence (channel FL5, excitation 488 nm, detection long pass 755 nm) (Fig. 1b,c). Pollen protoplasts demonstrate relatively low fluorescence in both channels. DiBAC_4_ fluorescence was measured in green part of spectrum (channel FL1, excitation 488 nm, detection 525/40 nm band pass).

### 3.6 ROS, antioxidants and inhibitors

We used freshly prepared 10 and 100 µM H_2_O_2_ from stabilized stock solution (Panreac) and catalase (Sigma); MnTMPP (200 µM) and catalase (100 U/ml) were added to exudate for 10 min to remove total ROS and selectively dismutate H_2_O_2_. LaCl_3_ (1 mM) and TEA (10 mM) were applied to protoplasts to inhibit Ca^2+^-permeable and K^+^-permeable channels as demonstrated previously (Breygina *et al*. 2016).

### 3.7 Protein extraction and trypsin digestion

Proteins were extracted by phenol extraction method (Faurobert *et al*. 2007). Plant tissue was homogenized in ice-cold extraction buffer (500 m Tris–HCl, pH 8.0, 50 mM EDTA, 700 mM sucrose, 100 mM КCl, 1 mM phenylmethylsulfonyl fluoride, 1 mM DTT), followed by 10 min incubation on ice. An equal volume of ice-cold Tris–HCl (pH 8.0)-saturated phenol was added, and the mixture was vortexed and incubated for 10 min with shaking. After centrifugation (10 min, 5500 × g, 4°C), the phenol phase was collected and re-extracted with extraction buffer. Proteins were precipitated from the final phenol phase with three volumes of ice-cold 0.1 M ammonium acetate in methanol overnight at −20°C. The pellets were rinsed with ice-cold 0.1 M ammonium acetate in methanol three times and with ice-cold acetone once and then dried. Resulting pellet was dissolved in 8 M urea, 2 M thiourea and 10 mM Tris. Proteins were quantified by Bradford protein assay (Bio-Rad, Hercules, CA USA). The 100 µg of proteins were reduced by 5 mM DTT for 30 min at 50°C and alkylated by 10 mM iodoacetamide for 20 min at room temperature. Proteins were dissolved in 40 mM ammonium bicarbonate and digested by 1 ug sequence-grade modified trypsin (Promega, Madison, WI, USA) at 37 °C overnight. Reaction was stopped by adding trifluoroacetic acid to 1%. 20 µg of each sample was desalted by Empore octadecyl C18 extraction disks (Supelco, USA) twice and then was dried in a vacuum concentrator.

### 3.8 LC-MS/MS analysis and peptide identification

Mass-spectrometry analysis was performed in three independent biological and three technical repeats. Reverse-phase chromatography was performed with an Ultimate 3000 Nano LC System (Thermo Fisher Scientific, Rockwell, IL, USA), which was coupled to a Q Exactive HF mass spectrometer (Q ExactiveTM HF Hybrid Quadrupole-OrbitrapTM Mass spectrometer, Thermo Fisher Scientific, USA). The peptides were separated in a 15-cm long C18 column with an inner diameter of 75 μm (Acclaim^®^ PepMap™ RSLC, Thermo Fisher Scientific, Rockwell, IL, USA). The peptides were eluted with a gradient from 5–35% buffer B (80% acetonitrile, 0.1% formic acid) over 65 min at a flow rate of 0.3 μL/min. Total run time including initial 10 min of column equilibration to buffer A (0.1% formic acid), then gradient from 5–35% buffer B over 65 min, 5 min to reach 99% buffer B, flushing 5 min with 99% buffer B and 5 min re-equilibration to buffer A amounted 90 min. MS analysis was performed at least in triplicate with a Q Exactive HF mass spectrometer (Q ExactiveTM HF Hybrid Quadrupole-OrbitrapTM Mass spectrometer, Thermo Fisher Scientific, Rockwell, IL, USA). Mass spectra were acquired at a resolution of 120,000 (MS) and 15,000 (MS/MS) in an m/z range of 300−1500 (MS) and 100– 2000 (MS/MS), correspondently. An isolation threshold of 100,000 counts was determined for precursor’s selection; up to top 20 precursors were chosen for fragmentation with high-energy collisional dissociation (HCD) at 29 NCE and 100 ms accumulation time. Precursors with a charged state of +1 were rejected and all measured precursors were excluded from measurement for 20 s. Other settings: charge exclusion-unassigned, 1, >6; peptide match – preferred; exclude isotopes – on; dynamic exclusion −20 s was enabled.

The raw mass-spectrometry data were processed using a freely available software suit, MaxQuant (v1.6.6.0). The MS/MS spectra were search against a database containing the protein sequences from Uniprot (73066 entries) and sequences from a database of common contaminant proteins (Tyanova, Temu, & Cox 2016). The false discovery rate (FDR) for peptide and protein identification was set to 0.01. The fixed modification (cysteine carbamidomethylation) and variable modification (methionine oxidation and N-terminal acetylation) were set for the search. All other parameters were left at default values. The MaxQuant’s LFQ algorithm which combines and adjusts peptide intensities into a protein intensity value was used for protein quantification.

The LFQ intensities of proteins from the MaxQuant analysis were imported in the freely available software Perseus (version 1.6.0.7) and transformed to logarithmic scale with base two (Tyanova, Temu, Sinitcyn, *et al*. 2016). Reverse database hits and contaminants were removed before performing two-way Student-t test and error correction (p value < 0.05) using the method of Benjamini–Hochberg was carried out. A minimum of 2 measurements per group was required. All those proteins that showed a fold-change of at least 1.2 and satisfied p < 0.05 were considered differentially expressed.

Gene ontology (GO) enrichment analysis was made using the Plant Transcriptional Regulatory Map online service (http://plantregmap.cbi.pku.edu.cn) (Jin *et al*. 2015, 2016).

### 3.9 Statistical analysis

Experiments were performed in four to seven independent replications. Results in the text and figures except the original pictures are presented as means ± standard errors. Significant difference was evaluated according to Student’s *t*-test. (\**p*< 0.05, \*\**p*< 0.01).

## 4 Results

### 4.1 Stigma exudate of tobacco contains ROS

Spectrofluorometrical measurements demonstrated a boost of DCF signal in exudate solution compared to PBS in which the washout was performed (used as baseline). The signal was strongly affected by MnTMPP – after the addition of H_2_O_2_ and O_2_^-**•**^ quencher fluorescence was severely reduced (Fig. 2). Thus, H_2_O_2_ and/or O_2_^-**•**^ are contained in stigma exudate in significant quantities.

**Figure 2.**
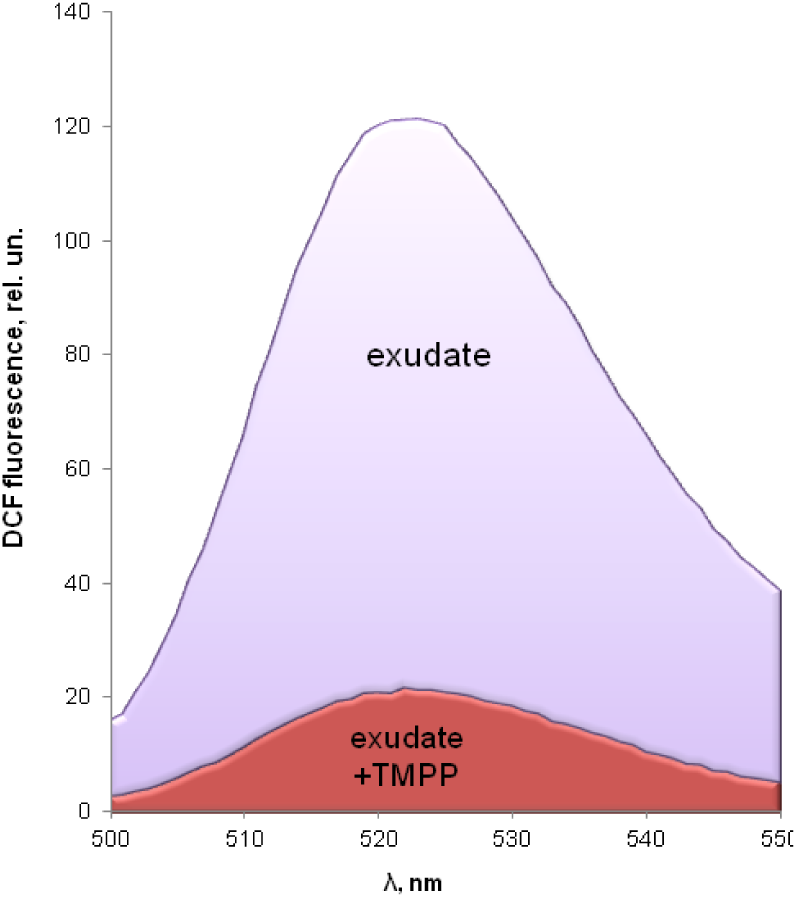
ROS detection is tobacco stigma exudate. Spectrum of DCFH fluorescence in the presence of stigma exudate before and after ROS quencher Mn-TMPP (200 µM) was added.

### 4.2 Stigma exudate and exogenous H_2_O_2_ affect membrane potential

We used MP mapping in growing pollen tubes to test MP as a complex parameter that can be affected by active substance produced by female tissues. In control pollen tubes, MP gradient was found, with the apical part depolarized compared to the shank (Fig. 3). The application of exudate to pollen suspension immediately causes hyperpolarization in apical and subapical parts of the tube (P<0.01 for 2 and 15 µm from the tip), in the basal part the difference was within the error. In treated tubes, the gradient persisted. As the measurement of MP gradient in single tubes is very laborious and impossible to perform for hundreds of tubes, we have designed a simple method of MP assessment which is fast and shows the distribution of the parameter in a large population – flow cytometry. We used protoplasts isolated from pollen tubes grown in the same conditions as those on which the effect of exudate was found, and the same exudate concentration. However, we used another optical method of MP assessment, and found the same effect as in pollen tubes.

**Figure 3.**
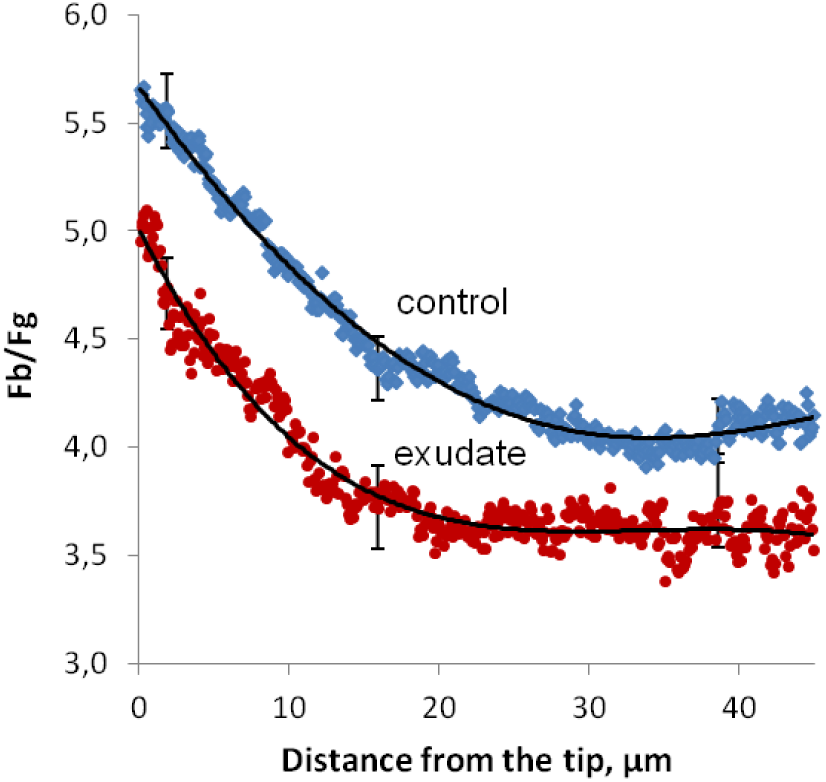
Stigma exudate causes hyperpolarization in pollen tubes. MP mapping in tobacco pollen tubes stained with Di-4-ANEPPS. Significant hyperpolarization can be seen in apical and subapical parts of tubes treated with stigma exudate (n=15) compared to control (n=31).

Population of protoplasts showed relatively stable DiBAC_4_(3) fluorescence which was strongly affected by an aliquot of stigma exudate. Shift of fluorescence towards lower reflects hyperpolarization of protoplasts (Fig. 4a). To test the system we applied NaN_3_ to cells and registered strong depolarization within 2 min (data not shown). H_2_O_2_ when added to protoplast suspension caused the same effect as exudate (Fig. 4b, e) indicating that H_2_O_2_ is the active substance in the exudate. 10 µM caused a modest hyperpolarization compared with exudate, while 100 µM caused a more pronounced effect (Fig. 4e), indicating that H_2_O_2_ concentration in the exudate is probably in this range. This assumption was further confirmed by experiments with catalase: 10 min pre-incubation of exudate with catalase (100 U/ml) completely abolished the effect on MP of protoplasts (Fig. 4c, d). The same abolishment was found in the presence of MnTMPP (200 µM) which quenched ROS in the exudate (Fig. 4d).

**Figure 4.**
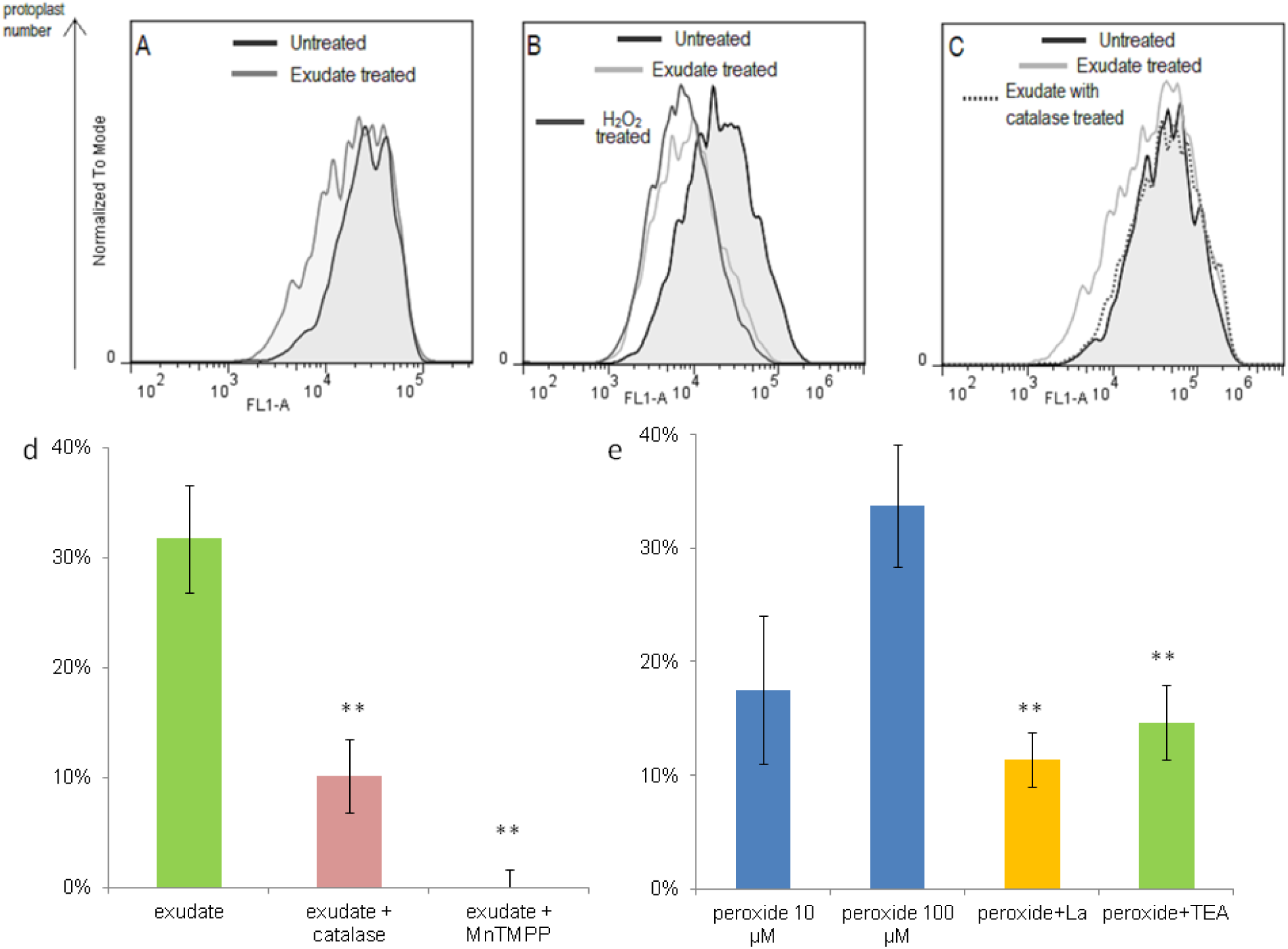
Hyperpolarization of pollen protoplasts in the presence of stigma exudate. a-c – distribution chart for protoplasts stained with DiBAC_4_(3) (examples of typical measurements). Decrease in fluorescence intensity reflects hyperpolarization in the presence of stigma exudate (a, b), H_2_O_2_ (b). After treatment with catalase stigma exudate lost the ability to cause hyperpolarization (c). d, e – mean hyperpolarization for all experiments (normalized to control) in the presence of exudate (d) without or with catalase/ROS quencher MnTMPP; (e) in the presence of H_2_O_2_ with or without cation channel blockers.

Hyperpolarization caused by 100 µM H_2_O_2_ was reduced almost 3 times in the presence of ion channel inhibitors – TEA and LaCl_3_ – thus indicating participation of K^+^- and Ca^2+^- conducting channels in the observed effect (Fig. 4e).

### 4.3 Pollen proteome is modified in the presence of ROS

We used label-free quantification approach to study pollen proteome changes after H2O2 treatment. Using mass-spectrometry analysis of three biological repeats, 1540 proteins were identified (Fig. 5a). To increase accuracy only proteins which were identified in at least two biological repeats by 2 or more unique peptides were analyzed; 733 proteins were identified in this case. Differently accumulated proteins (DAPs) were selected with significant changes (p-value < 0.05) and fold change above 1.2. A total of 20 DAPs was identified, 50 proteins were unique for peroxide-treated pollen and 4 for control samples (Fig. 5b, Table 1). Among DAPs, we found 11 proteins with increased abundance and 9 with reduced abundance. Gene ontology enrichment analysis for these groups was made, and the most impressive (and statistically relevant) results have been obtained for unique proteins in H2O2-treated pollen.

**Table 1.** Unique proteins in H_2_O_2_-treated pollen

| Protein ID | Description | Function (by similarity) |
| --- | --- | --- |
| A0A1S4APR3_TOBAC | luminal-binding protein 4-like | facilitates the assembly of multimeric protein complexes inside the ER |
| A0A1S4AUN8_TOBAC | NADH dehydrogenase [ubiquinone] iron-sulfur protein 7, mitochondrial-like | core subunit of the mitochondrial membrane respiratory chain NADH dehydrogenase |
| A0A1S3XTB8_TOBAC | heat shock cognate 70 kDa protein 2-like | stress response |
| A0A1S4D3H6_TOBAC | triosephosphate isomerase, | catalyzes the interconversion between DHAP and |
|  | cytosolic-like | G3P in glycolysis and gluconeogenesis |
| A0A1S3X6A4_TOBAC | pyruvate kinase | catalyzes the last step of glycolysis |
| A0A1S3WYM6_TOBAC | actin-101 | actin |
| A0A1S3YIE9_TOBAC | S-formylglutathione hydrolase | involved in the detoxification of formaldehyde |
| A0A1S4ALJ9_TOBAC | glucose-1-phosphate adenylyltransferase | plays a role in synthesis of starch. It catalyzes the synthesis of the activated glycosyl donor, ADP-glucose from Glc-1-P and ATP |
| A0A1S4AJF9_TOBAC | serine-tRNA ligase-like | catalyzes the attachment of serine to tRNA(Ser) |
| A0A1S3YDT5_TOBAC | At2g39795, uncharacterized protein, mitochondrial-like | positive regulation of mitochondrial translation |
| A0A1S4BBV8_TOBAC | ATP-dependent Clp protease proteolytic subunit | plays a major role in the degradation of misfolded proteins |
| A0A1S4A0H0_TOBAC | FAM10 family protein At4g22670-like | protein dimerization activity; chaperone cofactor-dependent protein refolding |
| W8R6T6_TOBAC Cys | cysteine synthase | cysteine biosynthesis from serine |
| A0A1S3XVM3_TOBAC | 20 kDa chaperonin, chloroplastic-like | co-chaperone |
| A0A1S3XU77_TOBAC | heat shock cognate protein 80-like | chaperone that promotes the maturation, structural maintenance and proper regulation of proteins |
| A0A1S4CRI6_TOBAC | 26S proteasome non-ATPase regulatory subunit 4 homolog isoform X2 | component of the proteasome |
| A0A1S4DMJ0_TOBAC | uncharacterized protein At2g34160-like | rRNA processing |
| A0A0M4JNG5_TOBAC | NADH dehydrogenase subunit 7 | core subunit of the mitochondrial membrane respiratory chain NADH dehydrogenase |
| A0A1S4C4G8_TOBAC | 60S ribosomal protein L38 | structural constituent of ribosome |
| A0A1S3ZJS3_TOBAC | NADP-dependent D-sorbitol-6-phosphate dehydrogenase-like | a key intermediate in the synthesis of sorbitol |
| A0A1S3XVZ1_TOBAC | carbonyl reductase [NADPH] 1-like | NADPH-dependent reductase with broad substrate specificity |
| A0A1S4B9Y3_TOBAC | uncharacterized protein At2g27730, mitochondrial-like | subunit of the mitochondrial respiratory chain complex I |
| A0A1S4AV83_TOBAC | leucine aminopeptidase 2, chloroplastic-like | involved in the processing and regular turnover of intracellular proteins |
| A0A1S4CJ13_TOBAC | pyrophosphate-fructose-6-phosphate 1-phosphotransferase subunit alpha | regulatory subunit of pyrophosphate--fructose 6-phosphate 1-phosphotransferase (glycolysis) |
| A0A1S3XST0_TOBAC | rho GDP-dissociation inhibitor 1-like | regulates the GDP/GTP exchange reaction of the Rho proteins |
| A0A1S3Y729_TOBAC | malic enzyme | component of the Citric acid cycle |
| A0A1S3Z2Y1_TOBAC | H/ACA ribonucleoprotein complex subunit 1-like isoform X2 | ribosome biogenesis, rRNA processing |
| A0A1S4BT67_TOBAC | UBP1-associated protein 2C-like | acts as component of a complex regulating the turnover of mRNAs in the nucleus |
| A0A1S4DNE8_TOBAC | peroxisomal fatty acid beta-oxidation multifunctional protein AIM1-like | is involved in the pathway fatty acid beta-oxidation, which is part of Lipid metabolism |
| A0A1S4CUN3_TOBAC | 2-dehydro-3-deoxyphosphooctonate aldolase 1 | lipopolysaccharide biosynthesis |
| A0A1S4B4V0_TOBAC | coatomer subunit zeta-1-like | protein transport from the ER, via the Golgi up to the trans Golgi network |
| A0A1S3Z1B3_TOBAC | pectinesterase | modification of cell walls via demethylesterification of cell wall pectin |
| A0A1S4AI50_TOBAC | mitochondrial-processing peptidase | required for maturation of the majority of |
|  | subunit alpha-like | mitochondrial precursor proteins |
| A0A1S3XD32_TOBAC | guanine nucleotide-binding protein subunit beta-like protein | component of ribosome |
| A0A1S4C023_TOBAC | DEAD-box ATP-dependent RNA helicase 56-like isoform X1 | involved in pre-mRNA splicing. Required for the export of mRNA out of the nucleus. |
| A0A1S3ZDB0_TOBAC | probable ADP-ribosylation factor GTPase-activating protein AGD8 | endoplasmic reticulum to Golgi vesicle-mediated transport |
| A0A1S3YBC0_TOBAC | COP9 signalosome complex subunit 5b-like | an essential regulator of the ubiquitin (Ubl) conjugation pathway |
| A0A1S3ZK66_TOBAC | dihydroxy-acid dehydratase, chloroplastic-like | amino-acid biosynthesis |
| A0A1S3ZZR4_TOBAC | ATP phosphoribosyltransferase 2, chloroplastic-like | amino-acid biosynthesis |
| Q9S820_TOBAC NTG | NTGP2 | small GTPase mediated signal transduction |
| A0A1S4DA91_TOBAC | translationally-controlled tumor protein homolog | involved in calcium binding and microtubule stabilization |
| A0A1S3YD94_TOBAC | mitochondrial import inner membrane translocase subunit TIM10-like | mitochondrial intermembrane chaperone |
| A0A1S4C7I8_TOBAC | β-galactosidase | hydrolysis of terminal non-reducing β-D-galactose residues (carbohydrate metabolic process) |
| A0A1S4CDD3_TOBAC | fumarylacetoacetase-like | amino-acid degradation |
| A0A1S4D3E9_TOBAC | ubiquitin carboxyl-terminal hydrolase 12-like isoform X2 | deubiquitinating enzyme |
| A0A1S4C2I0_TOBAC | 60S ribosomal protein L31-like | structural constituent of ribosome |
| A0A1S4ASH4_TOBAC | xylulose kinase-like | xylose metabolism |
| A0A1S3XAI9_TOBAC | ADP/ATP carrier protein, mitochondrial | transmembrane transport |

**Figure 5.**
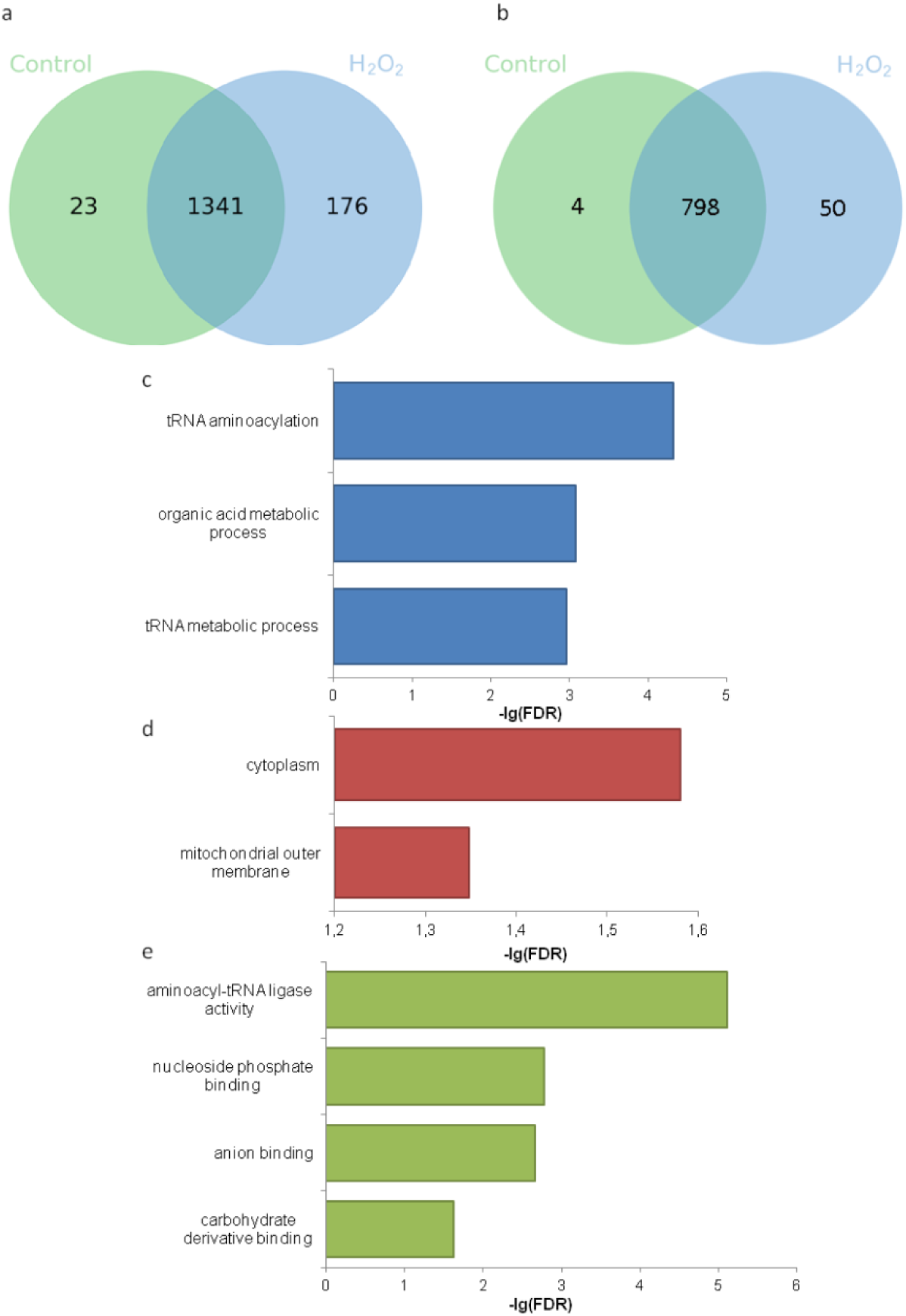
Venn diagram and GO enrichment of pollen proteins in the presence of 100 µM H_2_O_2_ (30 min treatment). a – Venn diagram of initially detected proteins; b - Venn diagram of proteins identified by at least 2 peptides; c-e – GO enrichment of unique proteins in treated pollen classified as biological processes (c); cellular component(d); molecular function (e).

Proteins were categorized into groups as biological processes, cellular components and molecular function. We found that among the unique proteins in the presence of 100 µM H_2_O_2_ three major groups attributed to biological processes were presented: tRNA metabolic process, organic acid metabolic process, tRNA aminoacylation (Fig. 5a). At least two of these groups constitute part of protein synthesis machinery, demonstrating its activation in the presence of ROS. Organic acid metabolism is an important part of synthetic processes such as rearrangement of carbohydrate skeletons and protein synthesis.

Cellular components mostly affected by 100 µM H_2_O_2_ are cytoplasm (translation machinery, metabolism) and mitochondrial outer membrane (translocation of proteins) (Fig. 5b) which again highlights the intensification of protein synthesis.

In molecular function category, carbohydrate derivative binding, nucleoside phosphate binding and aminoacyl-tRNA ligase activity were most abundant, indicating that these cellular activities were intensified.

To understand better functions of unique proteins in H_2_O_2_-treated pollen samples we divided them into several groups according to their functional annotation. The most abundant groups were Energetics (glycolysis, Crebs cycle, fatty acid oxidation, respiration), Metabolism (amino acid synthesis, rearrangement of carbohydrate skeletons, pentosophosphate cycle), Protein folding (chaperones), Translation and Regulatory proteins (Fig. 6). Energy and metabolism are probably activated in order to accelerate growth and activate the use of sucrose resources. Regulatory proteins and chaperones indicate an increase in protein synthesis, which agrees well with GO-enrichment.

**Figure 6.**
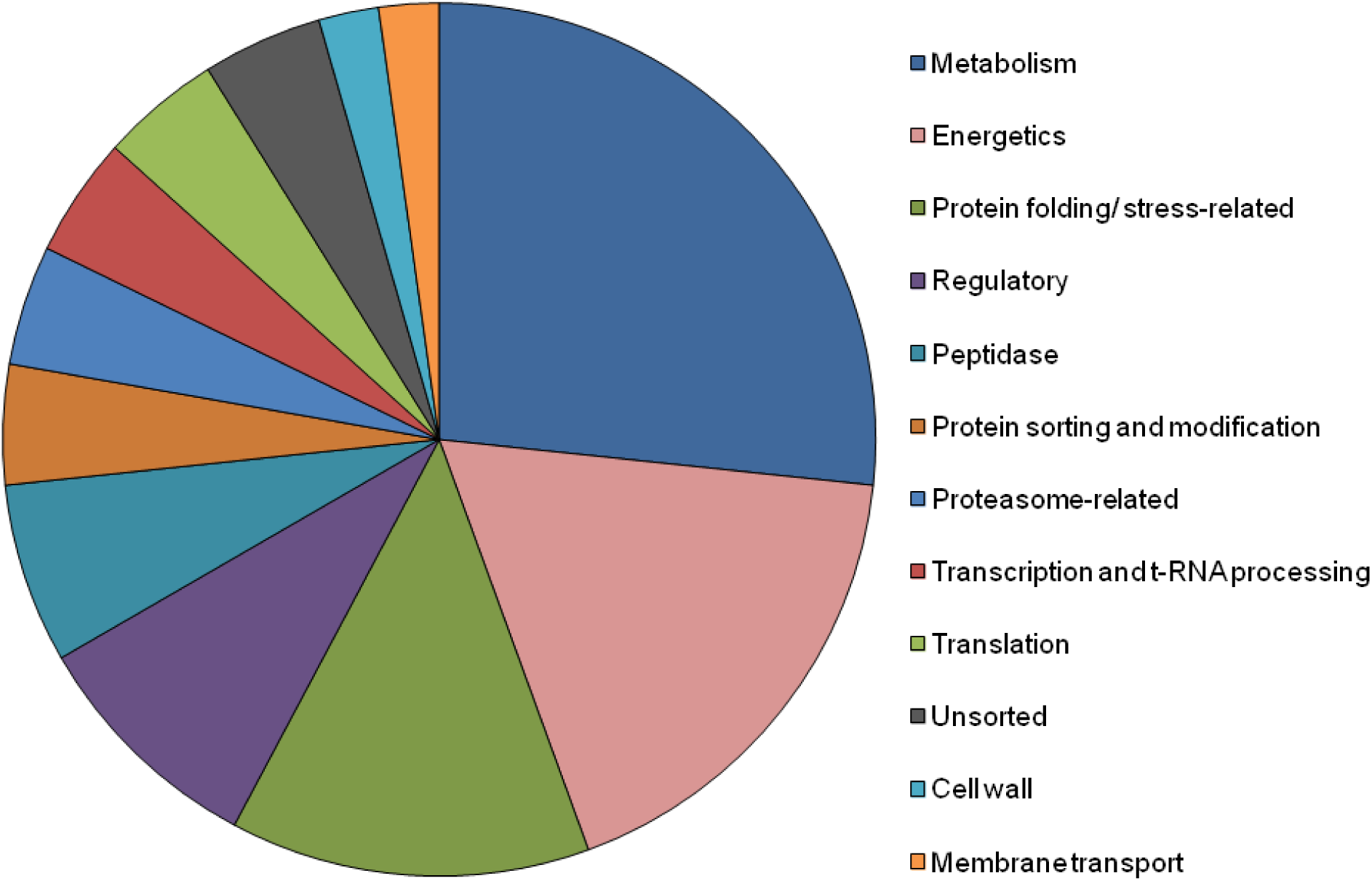
Functional classification of unique proteins in H_2_O_2_-treated pollen based on Uniprot database.

The list of unique proteins (Table 1) demonstrated the intensification of energy metabolism at all steps including glycolysis (2 proteins), Crebs cycle (1) and respiratory chain (3) as well as protein synthesis and translocation into mitochondria (2). The list includes some proteins that are important for polar growth regulation, for example, rho GDP-dissociation inhibitor 1-like (GDI-like) and NTGP2 (member of RAC/ROP GTPase gene family), which are components of RAC/ROP GTPase signaling. It is noteworthy that antioxidant proteins are missing in the list of unique proteins, thus, indicating normal physiological state of pollen cells which receive ROS signal. Three enzymes of amino-acid metabolism and five components of translation machinery were found in the list indicating the activation of translation. Several chaperons and associated proteins both cytolasmic and ER-localized are produced, including luminal-binding protein 4-like, which is the most abundant among the unique proteins. This emphasizes the growing need for protein folding, maturation, assembly of multimeric protein complexes, degradation of misfolded proteins etc.

Table 2 includes up- and down-regulated proteins in H_2_O_2_-treated pollen. Eleven up-regulated proteins represent the same main groups as unique proteins in H_2_O_2_-treated pollen: Energetics (PEP carboxylase), Metabolism (glucose and ribitol dehydrogenase, Ac-CoA carboxylase), Translation (eukaryotic initiation factor 4A-9, aspartate-tRNA ligase 2) and Protein folding (T-complex protein 1 subunit beta). We also found a highly overrepresented protein from polcalcin family of pollen allergens.

**Table 2.**
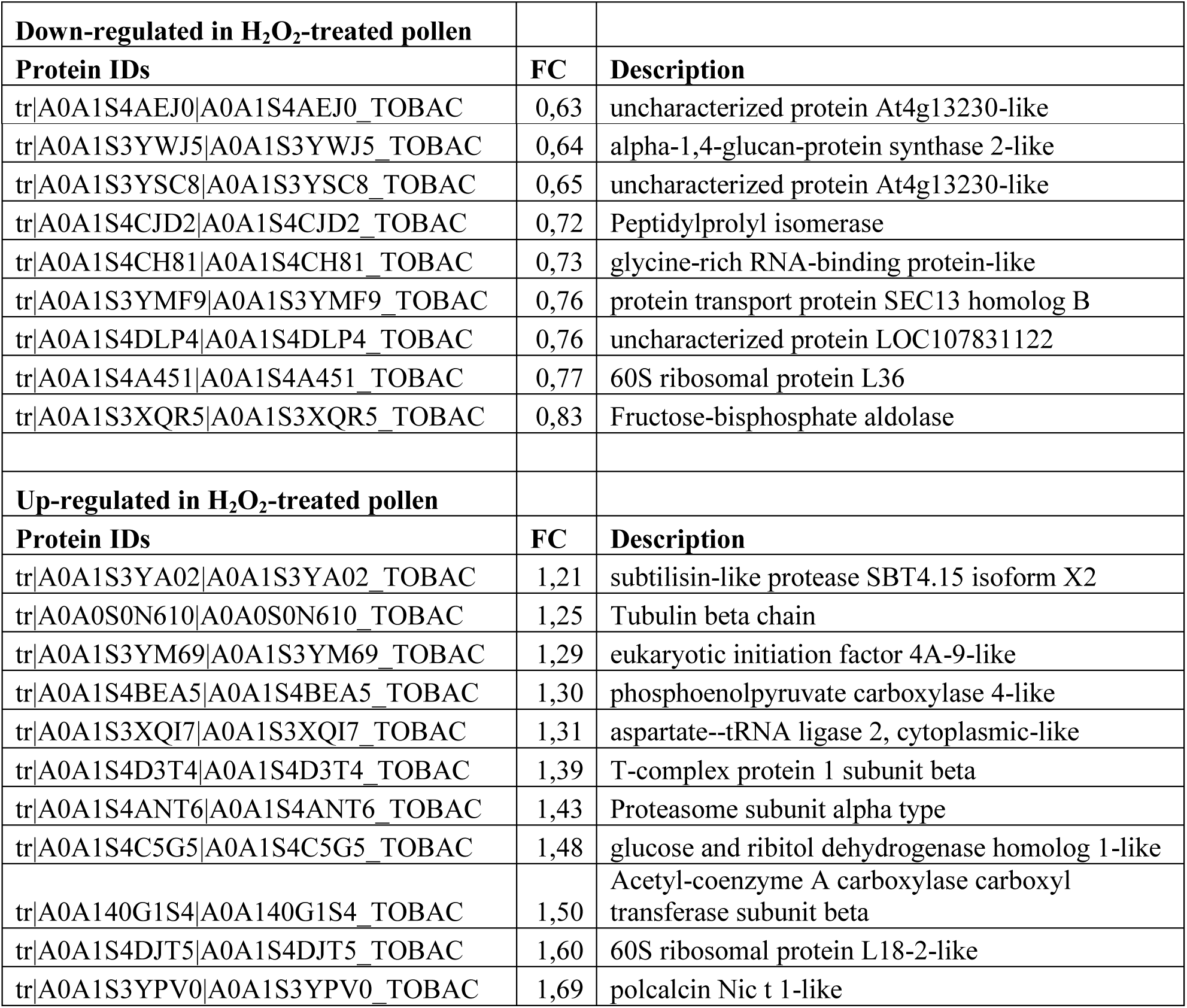
Up- and down-regulated proteins in H_2_O_2_-treated pollen

Among 9 down-regulated three proteins are uncharacterized, remaining 6 are presumably involved in: cell wall modification, mRNA transport, protein modification, RNA formation or processing during stress, ribosome structure. Aldolase catalyzes a reverse reaction that splits fructose 1,6-bisphosphate into triose phosphates. Anabolic pathways include the reverse reaction while the forward reaction is a part of glycolysis. The small number and significant proportion of uncharacterized proteins (1/3) do not allow accurate functional classification of down-regulated proteins but taking into account the ratio of up-regulated+unique proteins to down-regulated we can conclude that total protein synthesis is activated by ROS.

## 5. Discussion

The stigma exudate had been traditionally regarded as a reservoir of water, secondary metabolites, cell wall precursors and compounds that serve as energy supply for rapid pollen tube growth (Konar & Linskens 1966; Labarca *et al*. 1970), but recent studies revealed some active compounds, at least in several species (Rejón *et al*. 2014) that modulate pollen germination, such as calcium (Ge *et al*. 2009) and enzymes of catabolism and defense (Rejón *et al*. 2013). We found ROS in collected stigma exudate of tobacco, which agrees well with the previously reported fact of ROS synthesis and accumulation on stigmas of different species (McInnis *et al*. 2006a; Zafra *et al*. 2016), including sunflower (Sharma & Bhatla 2013), olive (Zafra *et al*. 2010), *Senecio squalidus* L. and *Arabidopsis thaliana* (McInnis *et al*. 2006b). ROS signaling in pollen germination has been assumed and discussed (Wudick & Feijo 2014) and putative mechanisms of ROS action on male gametophyte have been revealed *in vitro* (Wu *et al*. 2010; Breygina *et al*. 2016; Maksimov, Breygina, *et al*. 2016; Podolyan *et al*. 2019). In particular, exogenously added H_2_O_2_ caused plasma membrane hyperpolarization (Maksimov, Breygina, *et al*. 2016; Podolyan *et al*. 2019) in pollen tubes and protoplasts but the question remained whether this response appears in the presence of exudate or not. Here we demonstrate that hyperpolarization appears both in pollen tubes exhibiting normal MP gradient as reported earlier (Breygina *et al*. 2010) and in pollen protoplasts treated with stigma exudate, and is abolished after catalase treatment – thus, hyperpolarization is caused by H_2_O_2_ in the exudate and is a part of natural physiological crosstalk. The experimental system used can be considered as a useful model for assessment of membrane effects by flow cytometry. The suspension contains intact pollen grains and exines as well as pollen protoplasts, which can produce some antioxidant effects previously reported (žárský *et al*. 1987; Smirnova *et al*. 2012), which is quite close to the situation *in vivo*; however, this does not neutralize the effect of exudate or peroxide; obviously, the sensitivity to H_2_O_2_ is quite high.

The link between hyperpolarization and other effects of H_2_O_2_ previously found – activation of Ca^2+^- and K^+^-conducting channels (Wu *et al*. 2010; Breygina *et al*. 2016) – has been reported here by inhibitory analysis: blockage of these channels significantly reduced hyperpolarization induced by H_2_O_2_ thus thereby revealing the mechanism of hyperpolarization.

Unique proteins synthesized in the presence of H_2_O_2_ were found in a proteomic study. Proteomes of germinating pollen in angiosperms are believed to include 3 parts: presynthesized at maturation, translated during activation from presynthesized mRNA and transcripted/translated *de novo* (Holmes-Davis *et al*. 2005; Noir *et al*. 2005; Dai *et al*. 2006; Zou *et al*. 2009). ROS signal perceived for half an hour could affect the second and third groups, activating translation of mRNAs and/or transcription of genes involved in redox-response. We analyzed proteins that were absent in control but were detected in H_2_O_2_-treated pollen. The main functional groups of proteins were close to those involved in normal metabolism activation and polar growth start-up (Dai *et al*. 2006; Zou *et al*. 2009): metabolism, energetics, protein folding, signaling. We assume that H_2_O_2_ during long-term exposure drives pollen to accelerate preparation for growth but does not provoke synthesis of antioxidants, ion channels or other special proteins. Specific local response to ROS appears to function largely at the posttranslational level, for example, modifications of channels’ conductivity and subsequent amplification of ion currents (Wu *et al*. 2010; Breygina *et al*. 2016). These data are in good agreement with a previously established fact that 100 µM H_2_O_2_ increases germination efficiency of tobacco pollen (Smirnova *et al*. 2009b). Thus, this ROS can be regarded as a normal component of the environment for pollen on stigma, which induces germination in angiosperms. Interestingly, for gymnosperms this pattern is not characteristic (Maksimov *et al*. 2018).

The list of unique proteins, besides the main groups associated with metabolism, energetics, translation and protein folding, includes regulatory proteins RhoGDI1-like (homolog of GDP dissociation inhibitor 1) and NTGP2 (member of RAC/ROP GTPase gene family), which are required components of RAC/ROP GTPase signaling (Kost 2008). Small GTPases from the RAC/ROP family represent crucial regulators of pollen tube growth, coordinating multiple downstream signaling pathways, including the dynamics of F-actin in the apex and the tip-focused calcium gradient in the apical region (Kost 2008; Zhang & McCormick 2010). Nt-RhoGDI2 was found to be co-expressed with GTPase Nt-Rac5 in tobacco pollen tubes (Klahre *et al*. 2006), and the two proteins interact with each other in yeast two-hybrid assays (Klahre *et al*. 2006). Pollen NADPH-oxidase is presumably controlled by Rac/Rop GTPases, as in pollen tubes overexpressing wild-type NtRac5 and constitutively active NtRac5 elevated ROS level was detected, while overexpression of dominant-negative NtRac5 led to a decrease in ROS (Potocký *et al*. 2012). Thus, two members RAC/ROP GTPase signaling are produced during pollen germination in the redox-active environment and, in turn, can modulate ROS production in pollen tubes through a feedback regulation.

Up-regulated proteins exhibit the same pattern of functional classification as unique proteins demonstrating the activation of energy consumption and synthetic processes. Among up-regulated proteins the most overrepresented is a member of polcalcin family. Polcalcins are pollen-specific proteins containing two EF-hands highly conserved in a variety of dicot and monocot species (Rozwadowski *et al*. 1999). Polcalcins are mainly studied as strong human allergens (Magler *et al*. 2010), but immunolocalization in certain model species showed that they are abundant cytosolic low-*M*r proteins located at or near the pollen tube surface (Rozwadowski *et al*. 1999). Polcalcins bind calcium and undergo changes in conformation, thus, should display regulatory function and perhaps is involved in signal transduction pathways (Ledesma *et al*. 1998). Up-regulated by H_2_O_2_, polcalcin can act as a signal mediator between ROS, Ca^2+^ and downstream effectors during pollen germination, which may be the subject of further research.

